# Capturing the blue-light activated state of the Phot-LOV1 domain from *Chlamydomonas reinhardtii* using time-resolved serial synchrotron crystallography

**DOI:** 10.1101/2023.11.06.565770

**Authors:** Guillaume Gotthard, Sandra Mous, Tobias Weinert, Raiza Nara Antonelli Maia, Daniel James, Florian Dworkowski, Dardan Gashi, Antonia Furrer, Dmitry Ozerov, Ezequiel Panepucci, Meitian Wang, Gebhard F. X. Schertler, Joachim Heberle, Joerg Standfuss, Przemyslaw Nogly

## Abstract

Light-Oxygen-Voltage (LOV) domains are small photosensory flavoprotein modules that allow converting external stimuli (sunlight) into intracellular signals responsible for various cell behavior (*e*.*g.,* phototropism and chloroplast relocation). This ability relies on the light-induced formation of a covalent thioether adduct between a flavin chromophore and a reactive cysteine from the protein environment, which triggers a cascade of structural changes that results in the activation of a serine/threonine (Ser/Thr) kinase. Recent developments in time-resolved crystallography may allow the observation of the activation cascade of the LOV domain in real-time, which has been elusive.

In this study, we report a robust protocol for the production and stable delivery of microcrystals of the LOV domain of phototropin Phot-1 from *Chlamydomonas reinhardtii* (*Cr*PhotLOV1) with a high-viscosity injector for time-resolved serial synchrotron crystallography (TR-SSX). The detailed process covers all aspects, from sample optimization to the actual data collection process, which may serve as a guide for soluble protein preparation for TR-SSX. In addition, we show that the obtained crystals preserve the photoreactivity using infrared spectroscopy. Furthermore, the results of the TR-SSX experiment provide high-resolution insights into structural alterations of *Cr*PhotLOV1 from Δt = 2.5 ms up to Δt = 95 ms post-photoactivation, including resolving the geometry of the thioether adduct and the C-terminal region implicated in the signal transduction process.

## Introduction

Phototropin protein (phot) is a blue-light photoreceptor found in plants and algae that is responsible for the cellular response to light stimulation from the environment (sunlight) (Briggs *et al*., 2001). For example, in the green algae *Chlamydomonas reinhardtii* (*C. reinhardtii* or *Cr*), phot allows the light-dependent regulation of several molecular processes (*e*.*g*., phototaxis, sexual differentiation, photoprotection) and control of gene expression (Huang & Beck, 2003; Im *et al*., 2006; Trippens *et al*., 2012; Petroutsos *et al*., 2016). The *C. reinhardtii* phot protein consists of two successive photosensory protein modules, LOV1 and LOV2 domains, and a Ser/Thr kinase effector domain (Huang *et al*., 2002) (**Fig. 1a**). The LOV domains are connected to the kinase through linker sequences whose structural conformation is dependent on the signaling state of the associated LOV domain (Okajima *et al*., 2014; Nakasone *et al*., 2019; Henry *et al*., 2020). Thus, LOV domains can therefore be considered as natural molecular light switches and they have found many applications in optogenetics in recent years (Wu *et al*., 2009; Rao *et al*., 2013; Baarlink *et al*., 2013; Strickland *et al*., 2012; Niopek *et al*., 2014; Van Bergeijk *et al*., 2015; Wang *et al*., 2016).

**Fig. 1.**
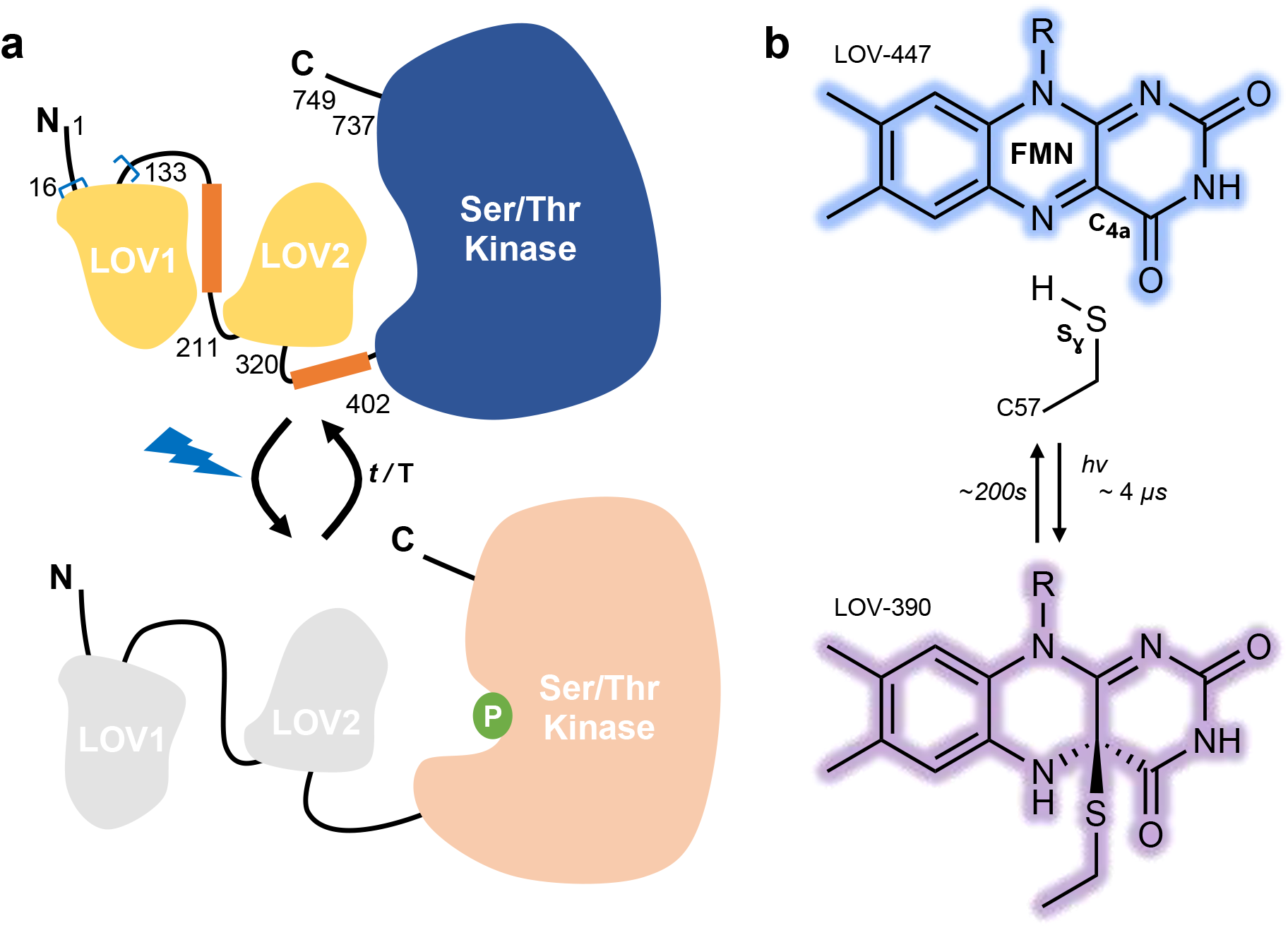
(**a**) Schematic representation of the architecture of the phototropin phot from *C. reinhardtii* showing the proposed mechanism of signal transduction. The investigated construct (amino acids [16-133]) is indicated between blue brackets. (**b**) Chemical structures of the dark state (LOV-447) and the light state (LOV-390).

LOV domains feature a flavin mononucleotide (FMN) chromophore with an absorption maximum at 447 nm under dark conditions (LOV-447) (**Fig. 1b**). Photoexcitation of the FMN chromophore induces the rapid formation of a triplet state on a nanosecond timescale, which then reacts with the thiol group of a cysteine residue from the protein to form a cysteinyl-FMN thioether covalent adduct after a few microseconds (Holzer *et al*., 2002; Kottke *et al*., 2003). This adduct exhibits an absorption maximum of around 390 nm (LOV-390). While activation is a fast process, the relaxation to the ground state is a thermal process occurring several orders of magnitude slower (∼ 200 s for *Cr*PhotLOV1) (Kasahara *et al*., 2002; Kottke *et al*., 2003).

The structural characterization of LOV debuted nearly two decades ago (Crosson & Moffat, 2001). However, the covalent adduct is particularly sensitive to specific X-ray radiation damage (Fedorov *et al*., 2003; Gotthard *et al*., 2019). Hence, first attempts to capture the light-adapted state were either performed at room temperature under continuous illumination where the continuous photoactivation leads to the accumulation of the adduct (Crosson & Moffat, 2002), or using the freeze-trapping method, after which several datasets are combined into a composite dataset of virtually lower accumulated X-ray dose (Fedorov *et al*., 2003). More recently, the progressive photoconversion from dark to the light-adapted state of *Arabidopsis thaliana* Phot2 LOV2 (*At*Phot2LOV2) domain was observed with a 63 ms time-resolution (Aumonier *et al*., 2020) following gradual population conversion within an expanding volume of crystal rather than direct time-resolved protein dynamics.

Pump-probe time-resolved (TR) serial femtosecond crystallography (TR-SFX) is a recent method that provided some of the most striking results on the dynamics of photoactive proteins on sub-milliseconds time scale (Tenboer *et al*., 2014; Kupitz, Basu *et al*., 2014; Barends *et al*., 2015; Nango *et al*., 2016; Nogly *et al*., 2018; Coquelle *et al*., 2018; Nass Kovacs *et al*., 2019; Skopintsev *et al*., 2020; Dods *et al*., 2021; Gruhl *et al*., 2023). On the other hand, its synchrotron counterpart, TR serial synchrotron crystallography (TR-SSX), has been successfully used to probe structural dynamics on a slower time scale (> ms) (Schulz *et al*., 2018; Weinert *et al*., 2019; Mehrabi *et al*., 2019). Both approaches are built on a similar principle and, considering the relatively higher accessibility of synchrotrons, offer powerful synergy (Mous *et al*., 2022). We report here the production of the *Cr*PhotLOV1 microcrystals (20 µm) necessary for an efficient extrusion and photoactivation and discuss the choice of a proper viscous matrix in which crystals are stable for the duration of the experiment. We show that the obtained crystals preserve the expected photoreactivity using infrared spectroscopy. Further, this work describes a TR-SSX experiment using a high-viscosity injector to study the *Cr*PhotLOV1 active state and provides a detailed view of LOV domain changes accompanying the active state formation. Our study serves as a case study and guidebook toward a successful TR-SSX experiment with soluble protein crystals using a high-viscosity injector.

## Methods

### Expression and purification

The genetic sequence coding for amino acids 16-133 of the LOV1 domain of *Chlamydomonas reinhardtii* phot1 protein was inserted into the pET16b expression plasmid between the restriction sites NdeI and XhoI. This allows the expression of a protein bearing an N-terminal His-tag. The expression was conducted in *Escherichia coli* BL21 DE3 by growing the cells in ZYP5052 auto-inducible medium (Studier, 2005) at 37°C until OD_600_ ∼ 1.0 and 17°C overnight. The protein was purified using nickel affinity chromatography with a 5 ml HisTrap HP column (GE Healthcare) followed by size exclusion chromatography on a HiLoad Superdex 75 16/600 column (GE Healthcare). Fractions corresponding to the protein were pooled and concentrated to 10 mg ml^-1^ for further crystallization.

### Crystallization

Limited proteolysis with trypsin removed the purification tag from the purified protein (adding 1:10 of 0.25 mg ml^-1^ trypsin solution). Crystallization screening was conducted to identify a condition producing a high density of microcrystals suitable for serial crystallography. The best condition consisted of 100 mM sodium cacodylate at pH 6.5 and 1.0 M sodium citrate dibasic trihydrate. Crystals of 10 - 30 µm in size appeared after one day using the sitting drop vapor diffusion with a 2:1 protein to precipitant ratio at 20 °C. Scaling up the crystallization and improving crystal size homogeneity were achieved in the batch crystallization method with seeding. Notably, crystals obtained during the first round of crystallization were used to prepare a seeding stock by crushing them with seeding beads (Hampton Research). Then the seeds were mixed with trypsin-digested protein (at 1:10 ratio). Finally, the mix was added dropwise in Eppendorf tubes containing the before-mentioned crystallization condition in a 2:1 ratio. Crystals with a size of 20 µm appeared the next day and slowly sedimented at the bottom of the Eppendorf tube.

### Sample preparation for serial synchrotron crystallography

A jetting solution of hydroxyethyl cellulose (23 % (w/v)) was prepared by dissolving dried cellulose in a solution containing the protein purification buffer and the crystallization condition in a 1:2 ratio. The cellulose mix was left to hydrate at room temperature until the medium became clear. Crystals were sedimented by centrifugation (800 x *g* for 1 min) and resuspended in the mother liquor for stabilization at the desired concentration. Resuspended crystals were inserted from the back of a Hamilton syringe and mixed in a 1:1 ratio with the hydrated viscous matrix using a 3-way syringe coupler (James *et al*., 2019).

### FTIR spectroscopy on CrPhotLOV1 crystals

Light-induced FTIR difference spectroscopy on protein crystals was performed essentially as described (Heberle *et al*., 1998). The FTIR difference spectrum in the 1800-1000 cm^-1^ range was recorded on a Vertex 80V spectrometer (Bruker) in attenuated total reflection (ATR) configuration (Nyquist *et al*., 2004), using a diamond ATR cell. For the 2620-2500 cm^-1^ range, the sample was sandwiched and sealed between two BaF_2_ windows and difference spectra were taken in transmission mode (Maia *et al*., 2021). In both configurations, crystals in mother liquor at pH 6.5 were kept in the dark for 300 s, followed by 10 s of illumination with a LED emitting at a center wavelength of 450 nm (∼10 mW cm^-2^). Overall, 3.200 light-dark difference spectra were recorded at a spectral resolution of 2 cm^-1^ and averaged.

### Cryogenic data collection at SLS

A LOV1 crystal was harvested and transferred to a cryoprotective solution consisting of the crystallization condition to which 20% glycerol was added. After equilibrating for 20 s, the crystal was fished from the cryoprotective solution and cryo-cooled in a 100 K nitrogen gas stream. Diffraction data were acquired at beamline X10SA (Swiss Light Source, Switzerland) with the fine slicing method by collecting 1800 images of 0.1° using a 73 × 16 µm^2^ beam width at a photon flux of 2 × 10^11^ photons s^-1^. Data were processed, scaled and merged using the XDS package (Kabsch, 2010). Data reduction statistics are presented in **Table 1**. Structure coordinates and structure factors have been deposited in the Protein Data Bank under the accession code 8KI8.

**Table 1.**
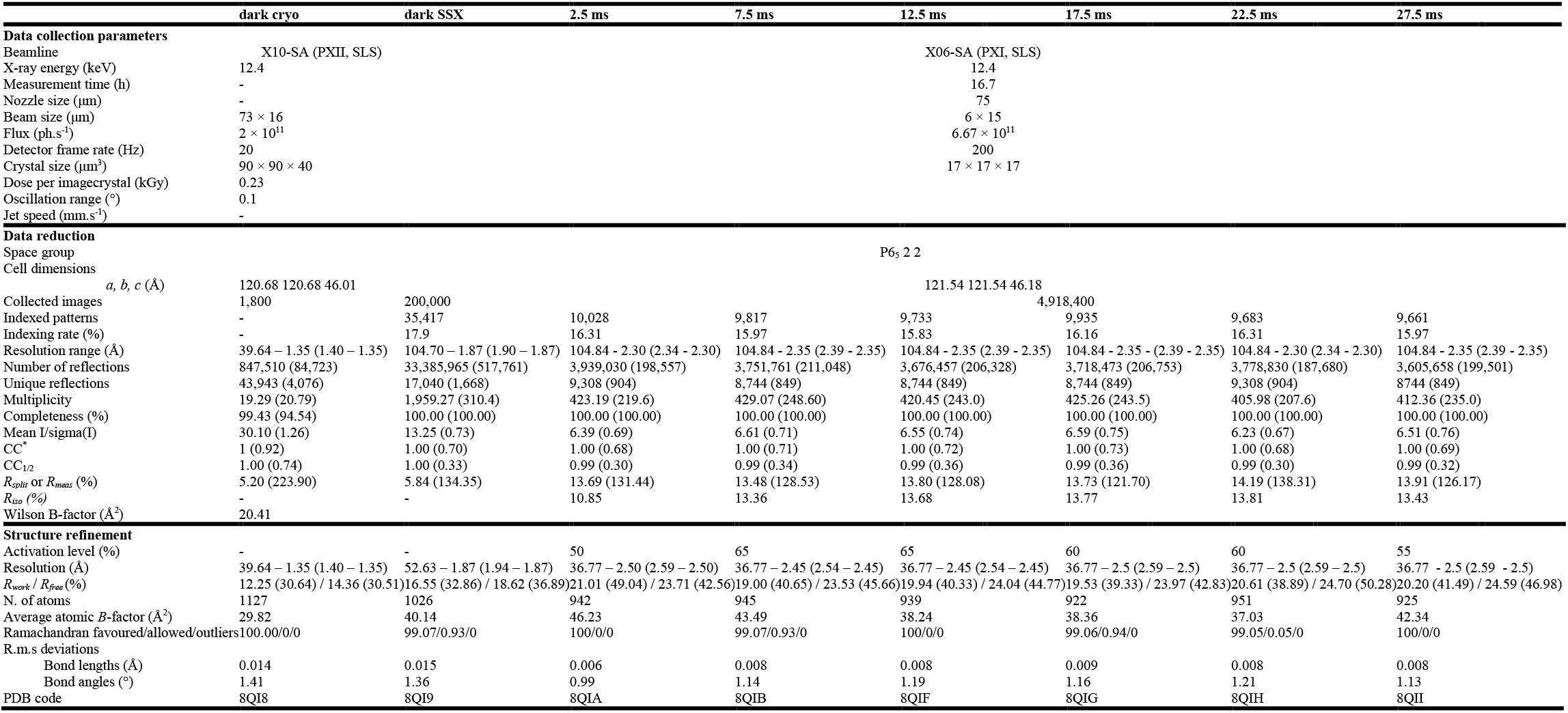
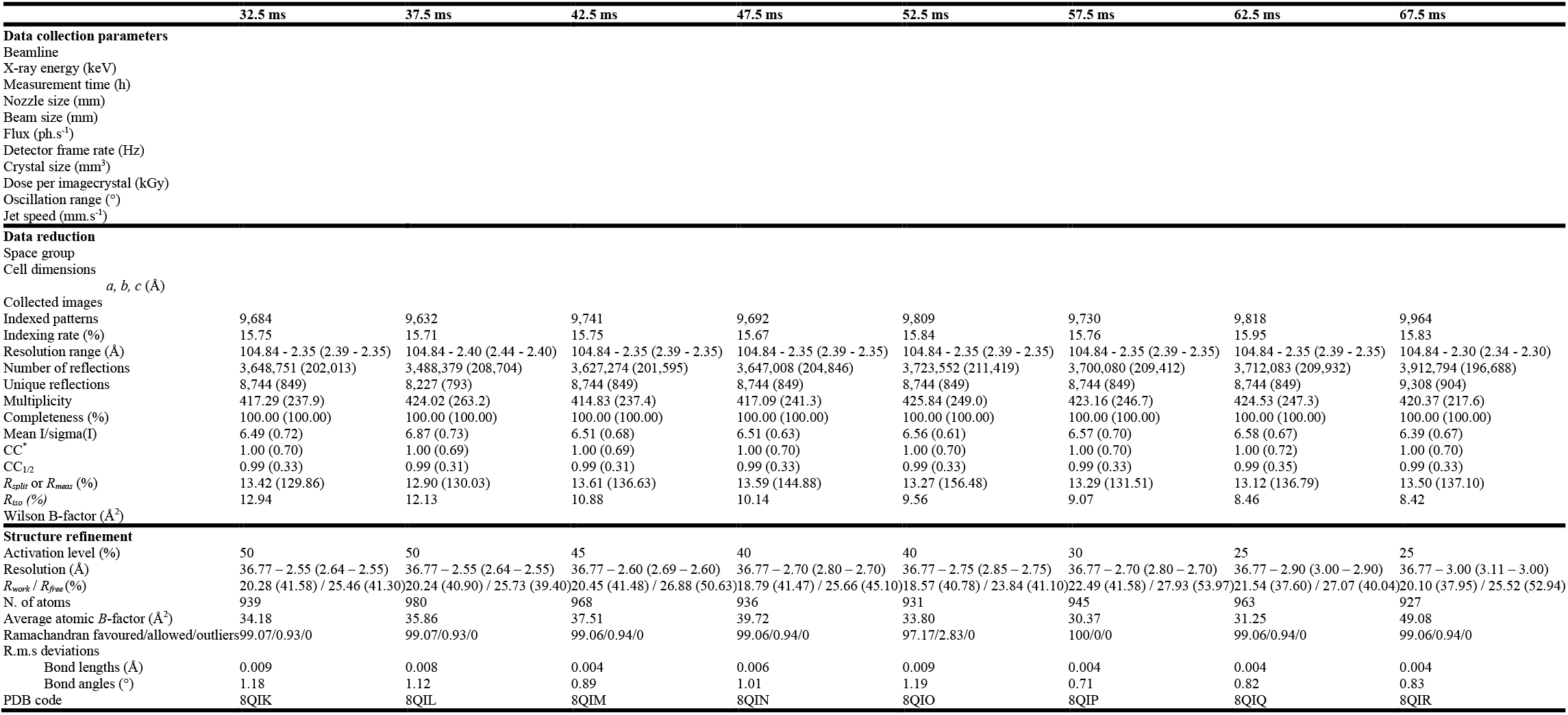
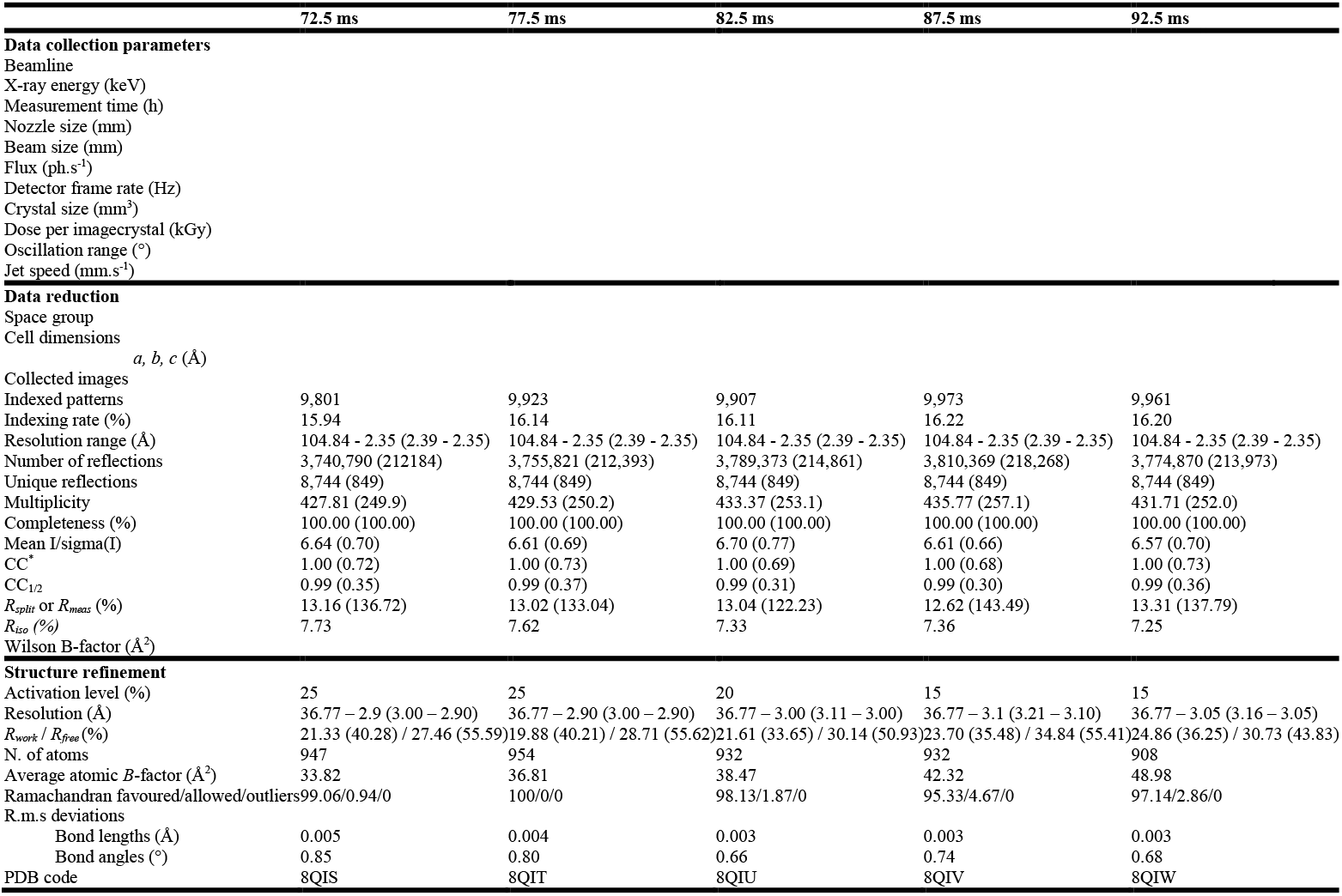
Data collection parameters, data reduction and refinement statistics.

### TR-SSX data collection and processing at SLS

Data were collected at beamline X06SA (Swiss Light Source, Switzerland) using the same setup as previously described (Weinert *et al*., 2017). Briefly, a stream of crystals was continuously extruded at the speed of 563 µm s^-1^ using a 75 µm nozzle onto the path of the continuous X-ray beam with a 15 × 6 µm^2^ beam width, 6.7 × 10^11^ photons s^-1^ flux and 12.4 keV photon energy. For the time-resolved experiment, a 5 ms light pulse of a 2.5 mW 488 nm pump laser diode was focused on a 104 × 170 µm^2^ 1/e^2^ spot (18 W cm^-2^) and synchronized with the detector trigger. The stability of the jet during the experiment was adjusted with a nitrogen gas sleeve. Diffraction patterns were collected using the central 4M region of an EIGER 16M detector recording at 200 Hz (as indicated in **Table 1**). The activation sequence was composed of one image collected with the laser diode on, followed by 79 images collected without illumination. This sequence was repeated 5 times, after which one activation sequence was skipped (**Supplementary Fig. S5**).

### Data processing

Serial data were processed using *CrystFEL* version 0.8.0 (White *et al*., 2012) after binning images corresponding to each time delay in the activation sequence (image 1 (Δt = 0 – 5 ms) will be labeled to Δt = 2.5 ms, image 2 (Δt = 5 – 10 ms) labeled Δt = 7.5 ms, etc., Δt up to Δt = 397.5 ms). Indexing and integration were performed with *indexamajig*, using the *xgandalf* (Gevorkov *et al*., 2019) and *mosflm* (Powell, 1999) algorithms, searching for peaks with a minimum signal-to-noise ratio of 4.2, using the unit cell parameters from the 100 K structure (*a* = 121.07 Å, *b* = 121.07 Å, *c* = 46.04 Å). Peak intensities were integrated using the *rings* method with *indexing radius* 4,5,9. Data were merged and scaled using the unity partiality model with a *partialator* with the unity partiality model and a *pushres* option of 1.8 nm^-1^. The resulting *hkl* files were converted into *mtz* with *ft2mz* from the *CCP*4 suite (Winn *et al*., 2011). A high-resolution cutoff was applied where CC_1/2_ was falling below 30%. Dataset statistics are reported in **Table 1**.

### Difference Fourier electron density maps

Fourier difference electron density maps were calculated using the *phenix.fobs_minus_fobs_map* program from the *Phenix* suite (Liebschner *et al*., 2019). A resolution cutoff of 2.1 Å and a sigma cutoff of 3.0 were applied and the multiscale option was used to calculate maps, subtracting dark data from the light data bins of interest as follows: *F* ^light^ – *F* ^dark^.

### Extrapolated electron density maps

The extrapolated structure factor amplitudes were calculated using a linear approximation (Genick *et al*., 1997) as follows: *F*_ext_ = [(*F*_obs_ ^light^ – *F*_obs_ ^dark^) / activated fraction] + *F*_obs_ ^dark^. The 2*F* – *F* maps calculated with phases of the dark state model showed distinct features in agreement with the *F*_obs_ ^light^ – *F*_obs_ ^dark^ Fourier difference maps. To infer activation levels, we calculated extrapolated maps with increasing steps of 5% of the activated fraction in F_ext_. This process continued until the dark state conformation features emerged on the Gln 120 side chain, at which point the activated fraction from the preceding step was utilized. The determined activation levels for different time bins are shown in **Fig. 6a**.

### Model building and refinement

Structures were solved using the molecular replacement method using *Phaser* (McCoy *et al*., 2007) and the structure coordinates of the LOV1 domain from *Chlamydomonas reinhardtii* (1N9L) solved by Fedorov and coworkers (Fedorov *et al*., 2003) as a search model. Several cycles of refining side chains and waters was performed using *Coot* (Emsley *et al*., 2010) and *Phenix* (Liebschner *et al*., 2019). Model representation and analysis were prepared with *Pymol* (http://pymol.org/). Coordinates and structure factors have been deposited in the Protein Data Bank with accession codes 8KI8 for the dark state structure obtained at cryogenic temperature, 8QI9 for the dark state structure obtained using serial crystallography at room temperature and 8QIA, 8QIB, 8QIF, 8QIG, 8QIH, 8QII, 8QIK, 8QIL, 8QIM, 8QIN, 8QIO, 8QIP, 8QIQ, 8QIR, 8QIS, 8QIT, 8QIU, 8QIV and 8QIW for the structures obtained by time-resolved crystallography at room temperature from 2.5 ms and to 92.5 ms after photoactivation (see Table 1).

## Results and discussion

### Sample preparation for a serial crystallography experiment

High-throughput serial crystallography experiments require the availability of microcrystals of the protein of interest in sufficient quantities (for an overview of suitable sample delivery methods, see Martiel *et al*. (Martiel *et al*., 2019) and Pearson & Mehrabi (Pearson & Mehrabi, 2020)). LOV domains yield crystals that can diffract to high resolution (**Supplementary Table S1**). Therefore, we first screened for crystallization conditions for *Cr*PhotLOV1 to identify spontaneously produced high density of micron-sized crystals in nanodrops (**Supplementary Fig. S1a**). Subsequently, the crystals were reproduced in 3 µl drops within 24-well plates, where various crystallization parameters, including protein-to-precipitant ratios and sample concentrations, were meticulously optimized. However, this approach yielded modest improvements as the differences between purification batches were difficult to control. To further improve the crystal quality, we applied limited proteolysis with trypsin as removing the expression tags was previously described to facilitate the crystallization of the homologous *At*Phot2LOV2 domain (Aumonier *et al*., 2020).

Ensuring the homogeneity of the crystalline sample is vital for obtaining optimal activation levels and promoting jetting stability in TR-SSX. Seeding can be employed to control the nucleation and the number of crystals, directly influencing the crystal size and the length of the crystallization experiment. The ratio between diffraction patterns and the total number of images recorded, commonly referred to as the hit-rate, is a vital parameter to consider. The crystal density of the sample determines the hit-rate during the SSX experiment and, thus, the efficiency of the data collection in the available time. Consequently, finely controlling crystal density would allow to further optimize the hit-rate in the serial experiment. We could readily generate crystal micro-seeds stock by crushing macrocrystals using a tissue grinder and resuspending them in the crystallization solution. This micro-seeds solution can then be employed to initiate crystallization in tubes via the micro-batch method (Kupitz, Grotjohann *et al*., 2014), thereby facilitating the growth of high-quality crystals for further analysis. Crystal size could be controlled by adjusting the volume of seeds (with a higher volume of seeds reducing the average crystal size) and length of crystallization (stopping the crystallization early allows obtaining smaller-sized crystals; **Supplementary Fig. S1b**). Overall, the crystallization process could typically be halted after one day through centrifugation, enabling the supernatant to be repurposed for an additional cycle of batch crystallization by incorporating new seeds. This method facilitated the generation of 5 µl of highly concentrated protein crystals suspension (approximately 5⋅10^6^ crystals ml^-1^) from a milligram of protein, featuring an average crystal size of 20 µm, which were well-suited for time-resolved serial synchrotron crystallography (TR-SSX) experiments.

### Choice of a carrier matrix for viscous injection

The lipidic cubic phase (LCP) injector, or high-viscosity extruder (HVE) (Weierstall *et al*., 2014, Botha *et al*., 2015), and high-viscosity cartridge-type (HVC) injector (Shimazu *et al*., 2019) are known for their extremely low flow rates (0.1-1 µl min^-1^) that result in low stream velocities (28-281 µm s^-1^). As a result, they drastically reduce sample consumption and enable efficient serial data collection at synchrotrons (Botha *et al*., 2015; Nogly *et al*., 2015). This delivery method is particularly suitable for membrane protein crystals (Jaeger *et al*., 2016) grown in the LCP mesophase (Landau & Rosenbusch, 1996) and has been shown effective for TR-SFX experiments (Nogly *et al*., 2016) and TR-SSX (Weinert *et al*., 2019). However, the viscosity of soluble protein crystals dispersed in precipitant solution is generally too low for high-viscosity delivery methods, necessitating the adjustment of the crystalline sample with the addition of grease or polymers (Nam, 2019).

At the beginning of the project, various crystal carrier media were evaluated for their efficacy. We first assessed if the crystals survived mixing with the carrier matrix by a visual inspection under the microscope. *Cr*PhotLOV1 microcrystals (**Fig. 2a**) dissolved rapidly upon mixing with monoolein or superlube grease (**Fig. 2b** **& 2c**, respectively). We identified polyethylene oxide (PEO) (Martin-Garcia *et al*., 2017) and hydroxyethyl cellulose (HEC) (Sugahara *et al*., 2017) as potential candidates. We then assessed the jetting properties of PEO and HEC by conducting a jetting experiment on an off-line setup consisting in an LCP-injector and a high speed camera allowing to observe the jet. Under our experimental conditions, PEO displayed unsatisfactory jetting properties as the jet diameter expanded after extrusion from the nozzle (data not shown). This high-viscosity matrix was therefore excluded as its expansion could potentially impact diffraction properties, induce unit cell expansion, and increase the path length of the activating light pulse. Eventually, we identified HEC as the optimal carrier matrix for *Cr*PhotLOV1 microcrystals. HEC was previously shown to be suitable for TR-SFX (Tosha *et al*., 2017; Wranik *et al*., 2023). Despite its moderate absorption in the UV spectrum, HEC is transparent at the excitation wavelength of 470 nm (Demina *et al*., 2020) used in our TR-SSX experiment. A highly concentrated crystalline protein sample was prepared for extrusion by gently mixing it with the rehydrated HEC matrix in Hamilton syringes using a three-way coupler (James *et al*., 2019). Visual inspection of the sample embedded in the HEC matrix indicated that the crystal integrity was maintained (**Fig. 2d**). Thus, HEC enabled the extrusion of 17 × 17 × 17 µm ± 4.3 µm crystals through the injector with a nozzle of 75 µm inner diameter, resulting in a stable jet with a stream velocity of 563 µm s^-1^ (**Fig. 2f**).

**Fig. 2.**
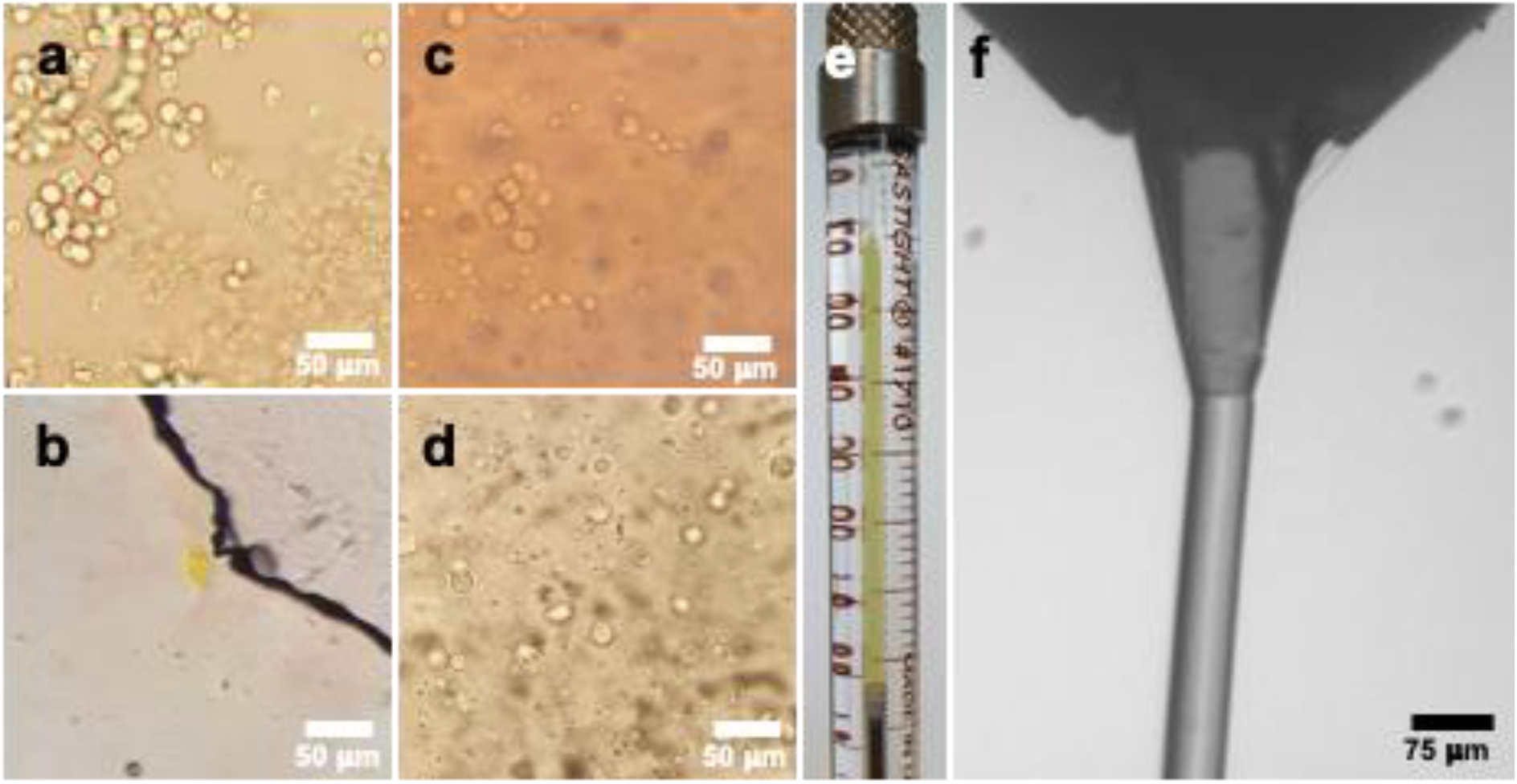
(**a**) LOV1 microcrystals in their crystallization solution, (**b)** after mixing with monoolein to prepare an LCP phase, (**c**) after mixing with superlube grease, and (**d**) after mixing with HEC. (**e**) Hamilton syringe containing LOV1 crystals mixed with HEC. (**f**) Close-up on the nozzle of the jet showing a stable extrusion with the HEC condition.

### Structure determination and refinement of the dark state at cryogenic temperature

To serve as a control experiment, we determined the dark state structure of *Cr*PhotLOV1 at cryogenic temperature (CT) from a single crystal (**Table 1**). Despite crystallizing under different conditions from those reported by Fedorov and colleagues (Fedorov *et al*., 2003), the crystals belonged to the same P6_5_ 2 2 space group, and diffraction data extended to 1.35 Å resolution - an improvement of 0.55 Å over the previously deposited dark state structure (PDB ID 1N9L). The recorded dark state structure superimposed well with the deposited structure, showing a root-mean-square deviation (RMSD) of 0.15 Å (measured on the backbone Cα over 104 residues). However, compared to the previously published structure, we observed that the Arg74 side chain had rearranged (Chi3 57° to 4°) as it accommodated a altered rotamer of the flavin phosphoribityl tail (**Supplementary Fig. S2**). This variation in the Arg74 and flavin tail conformations may have arisen from differences in the crystallization conditions and is not believed to influence the activation mechanism. The significant improvement in spatial resolution also allowed us to model Leu34, Val103, Ile73 and Cys32 residues surrounding the flavin in alternate conformations (**Supplementary Fig. S2a**), revealing system equilibrium dynamics and several water molecules coordinating the phosphoribityl tail and the phosphate group (**Supplementary Fig. S2b**).

### Dark-state structure at room temperature

Using the previously described setup (Weinert et al., 2017) and the LCP injector at the SLS beamline X06SA (PXI), we performed an SSX experiment with *Cr*PhotLOV1 crystals embedded in HEC. We collected 200,000 images in approximately 16.7 min, resulting in a sample consumption of 2.5 µl at a flow rate of 151 nl min^-1^ (**Table 1**). Of the 200,000 images, 35,871 diffraction patterns were successfully indexed and integrated, corresponding to an indexing rate of 17.9%. These patterns were merged to yield a dataset with a resolution of 1.87 Å, completeness of 100%, and a CC1/2 of 0.33 in the highest resolution shell (**Table 1**).

As expected from the cryogenic temperature characterization, the *Cr*PhotLOV1 crystals belonged to the P6_5_ 2 2 space group. We used the model coordinates of the CT dark state structure to calculate initial phases and then manually adjusted them with *Coot* before refining them with *Phenix*. Overall, the electron density was of excellent quality and enabled us to observe variations in the positions of residue side chains (with an RMSD of 0.189 Å between the dark state at CT and room temperature). The reactive cysteine (Cys57) exhibited two alternate conformations, as observed at CT, but the variation of the 2F_o_-Fc map contour at room temperature clearly indicated a change in the distribution of each conformation (**Fig. 3a** and **3b**, respectively). We thus refined the occupancy of cysteine using *Phenix* for both temperatures. Conformation A, in which the S_γ_ atom of Cys57 is 3.5 Å from the C4a of FMN, was equally present at room temperature along with conformation B (i.e., 0.50 and 0.50 for A and B conformations, respectively), in which the Sγ atom of Cys57 is 4.4 Å from the C4a of FMN.

**Fig. 3.**
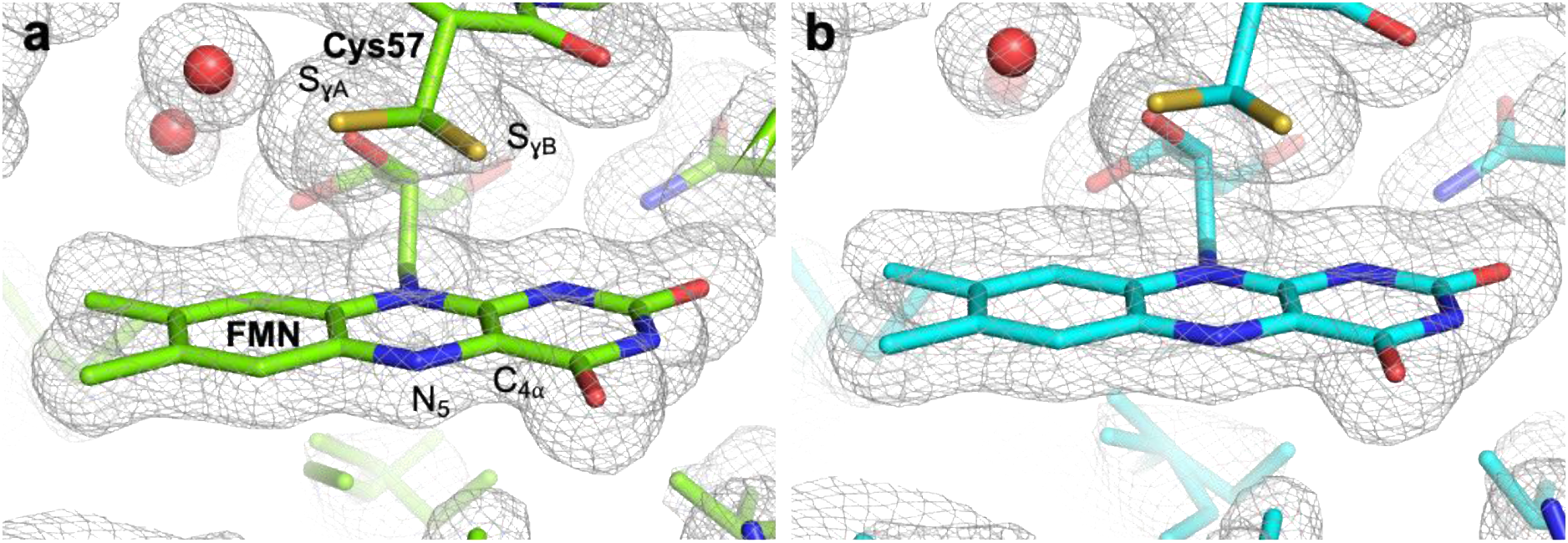
2*F*_obs_ – *F*_calc_ electron density maps contoured at 1 σ level around the FMN and the reactive Cys57 in (**a**) crystals collected at CT with the oscillation method, and (**b**) the SSX dataset collected at RT.

However, at cryogenic temperature, conformation A is favored (with an occupancy of 0.70 compared to 0.30 for conformation B). This observation is consistent with previous spectroscopic studies on the homologous LOV2 domain from *Adiantum neochrome* 1, which showed that conformation A is favored at low temperatures while adduct formation is more efficient with conformation B (Sato *et al*., 2007). The natural fluctuations between the different cysteine conformations occurring more frequently at physiological temperatures could potentially play a role in the recruiting process for the formation of the covalent adduct.

### Cr*PhotLOV1 is active in its crystalline form*

To investigate whether *Cr*PhotLOV1 was reactive in our crystals prior to the TR-SSX experiment, we recorded a light-induced Fourier-transformed infrared (FTIR) difference spectrum on microcrystals. FTIR allows probing of light-induced changes in the vibrational modes of the FMN and protein that occur upon light excitation. In the difference spectrum shown in **Fig. 4a**, negative bands are related to vibrations of the dark-state *Cr*PhotLOV1 that change upon photoconversion to the adduct state, which is characterized by positive bands. The difference spectrum of crystalline *Cr*PhotLOV1 is very similar to that of *Cr*PhotLOV1 in solution (Ataka *et al*., 2003), except for alterations in the amplitudes that are caused by the anisotropic polarization conditions in attenuated total reflection (ATR) spectroscopy, which preferentially enhance some vibrational bands of the crystalline protein lattice. Light-induced adduct formation involves proton transfer from Cys57 to N_5_ of FMN, and the terminal sulfur atom forms a covalent bond with C4a of FMN. The negative band at 2568 cm^-1^ indicates the deprotonation of the thiol S-H of Cys57 (**Fig. 4b**), which is very similar to *Cr*PhotLOV1 in solution (Ataka *et al*., 2003). The vibrational band at 1711 cm^-1^ has been assigned to the stretching vibration of C_4_=O in dark-state *Cr*PhotLOV1 (Swartz *et al*., 2002; Ataka *et al*., 2003; Iwata *et al*., 2006). The C_4_=O bond gains strength upon the formation of the C4a–S adduct, as reflected by the frequency upshift to 1724 cm^-1^ (**Fig. 4a**). The other large difference bands are indicative for the light-induced conversion of planar oxidized flavin to the thioadduct with nearby Cys57. These results collectively indicate that *Cr*PhotLOV1 in the crystalline state is active and forms a covalent adduct under the crystallization conditions used for the TR-SSX experiment.

**Fig. 4.**
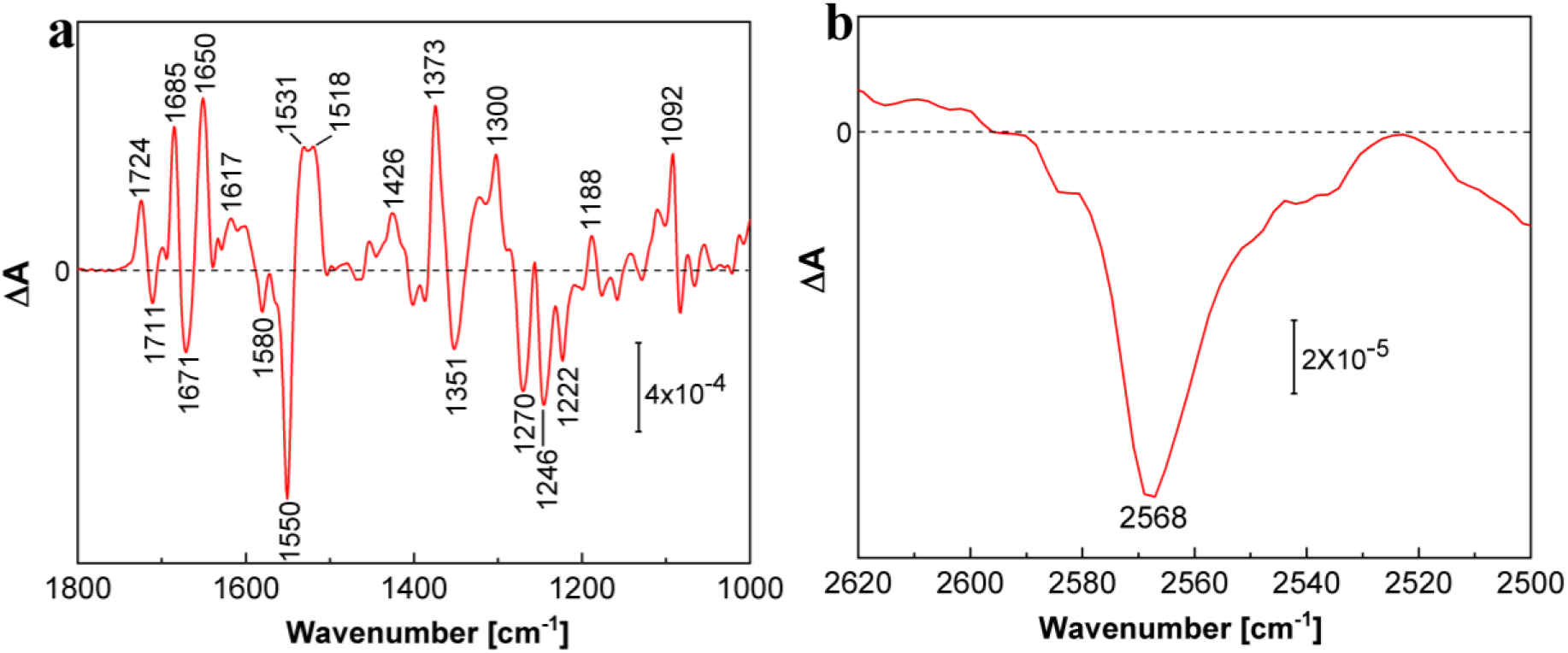
(**a**) Light-induced FTIR difference spectrum of *Cr*PhotLOV1 crystals. (**b**) Vibrational band of the S-H stretching vibration of Cys57. Crystalline *Cr*PhotLOV1 samples were photoactivated by a LED emitting at 450 nm.

### Structure determination of photoactivated states

To elucidate the light-induced structural changes occurring within the millisecond time domain, we employed pump-probe SSX. The experimental setup remained consistent with the previously described configuration (Weinert *et al*., 2019). In this approach, a delay generator synchronized data collection with a laser diode, as illustrated in **Supplementary Fig. S4**. During the experiment, LOV microcrystals were exposed to focused 488 nm laser diode light for 5 ms at the X-ray intersection region. Concurrently, the photocycle was probed by collecting 80 consecutive 5 ms frames, as depicted in **Supplementary Figure S5**. A total of 4,918,400 frames (61,480 per delay) were acquired over 6.8 hours, corresponding to a sample consumption of 62 µl (or 3.8 mg of protein) at a flow rate of 150 nl min^-1^. Of these images, 833,583 patterns were successfully indexed and integrated, resulting in an indexing rate of 16.9%. According to our data collection scheme, the first image in each sequence represents a time delay of 0 – 5 ms (Δt = 2.5 ms), with subsequent images corresponding to 5 – 10 ms (Δt = 7.5 ms) and so on, up to Δt = 397.5 ms. Images within each time delay bin were processed as separate datasets. Comprehensive statistics for the collected datasets are provided in **Table 1**.

### Addressing radiation damage concerns

The possibility of specific radiation damage (Holton, 2009; Garman & Weik, 2017), defined as site-specific alterations to protein structures or chemical bonds due to the ionizing effect of X-ray beams, was investigated. This type of damage affecting the covalent thioether adducts has been previously reported in multiple studies involving LOV proteins (Fedorov *et al*., 2003; Halavaty & Moffat, 2007; Zoltowski *et al*., 2007; Gotthard *et al*., 2019). Utilizing RADDOSE-3D (Zeldin *et al*., 2013), we calculated the accumulated dose per shot to be 15 kGy, considering a 50% overlap in crystal volume exposed to the X-ray between consecutive shots. This overlap occurred as the crystal translated by 3 µm per frame while the vertical beam dimension spanned 6 µm. Notably, this dose is approximately three times lower than the reported τ_1/2_ value of 49 kGy at room temperature (RT) observed in the homologous *At*Phot2LOV2 domain (Gotthard *et al*., 2019). The 49 kGy dose was delivered in a carefully devised low-dose data collection strategy, preventing any apparent signs of site-specific damage to the sensitive covalent adduct. Consequently, the light-activated state structures presented in the current study are likely to be predominantly unaffected by specific radiation damage, which would otherwise manifest through the reduction of the adduct, resulting in a dark state-like geometry.

### Examining activation levels in illuminated crystals

Structural changes can be examined through two distinct types of electron density maps: 1) Fourier-difference electron density maps (*F*_obs_ ^light^ – *F*_obs_ ^dark^), which involve using diffraction data collected without illumination as the dark reference and subsequently subtracting it from the data collected post-light exposure; 2) extrapolated maps, which facilitate the selective modeling of active state conformations by eliminating the dark state’s contribution to structure factor amplitudes (Genick *et al*., 1997). In the latter approach, the activation level of a map is determined by calculating and comparing extrapolated maps at varying activated fractions. The active state level is reduced until specific features corresponding to the dark state model (*e.g*., the dark state conformation of Gln120) are no longer present in the 2*F*_ext_ – *F*_calc_ electron density map. Intriguingly, our illumination conditions enabled the attainment of activation levels ranging from 65% (at Δ*t* = 7.5 ms) to 15% (at Δ*t* = 87.5 ms; **Fig. 6a**).

The high activation level may result from the relatively brief delay in adduct formation (∼ 4 µs) relative to the pump light pulse duration (5 ms), providing non-reacting species with multiple opportunities to react, and the remarkable stability of the Cysteinyl-FMN adduct. The excellent quality of the resulting extrapolated electron density maps facilitated the modeling of structural changes occurring post-light activation (Δ*t* = 2.5 - 92.5 ms; **Supplementary Fig. S6**).

### Analysis of light-induced structural changes

Fourier difference electron density maps reveal several positive (indicating incoming atoms) and negative (signifying outgoing atoms) peaks located around FMN (**Fig. 5a**). At 2.5 ms post-light activation, the most prominent features include a 15.8 σ peak located between Cys57 and C4a of FMN, along with a -7.5 σ peak on conformation A of Cys57. These observations are in line with the light-induced formation of the thioether covalent adduct (Crosson & Moffat, 2002; Halavaty & Moffat, 2007; Möglich & Moffat, 2007). The immediate structural consequences involve sp^3^ hybridization of the C4a atom, characterized by a -6.0 σ peak beneath the flavin plane and a 4.0° tilt of the isoalloxazine ring accompanied by a 4.4 σ positive density peak above the plane. In addition to covalent adduct formation, Gln120 has been proposed to participate in signal propagation (Iuliano *et al*., 2020). This key residue also displays strong features in the difference maps (8.2 σ (3rd most intense peak); -4.5 σ). In the resting state, the nitrogen atom of the Gln120 amide group forms a hydrogen bond with N_5_ of FMN. Refining the structure using extrapolated data enables the placement of the amide group’s oxygen atom near the strong positive peak, which, along with more consistent refined B-factors, indicate that the Gln120 amide rotates after the expected protonation of the N_5_ atom of FMN. Consequently, in the light-activated state, the oxygen atom of the Gln120 amide forms an H-bond with the N_5_ atom of the FMN chromophore (3.6 Å; **Fig. 5b**). Another result of Gln120 rotation is the weakened interaction with Thr21, transitioning from a strong hydrogen bond interaction with the Gln120 oxygen at 2.7 Å to an asymmetric hydrogen bond interaction with the nitrogen at 3.2 Å. The attenuation of interactions between the N-terminal and C-terminal regions may influence protein dynamics and contribute to signal transduction, as suggested for *As*LOV2 (Iuliano *et al*., 2020). This effect could destabilize the linker sequence to the LOV2 domain, subsequently releasing the kinase from its inactive form (Peter *et al*., 2010; Henry *et al*., 2020). Several other residues exhibit prominent features in the difference density map. In particular, Leu34, characterized by a pair of positive and negative peaks of 5.2 σ and -4.1 σ, moves towards the space vacated by the alternate conformation of Cys57 following adduct formation. This observation has also been reported in *A*tPhot2LOV2 (Aumonier *et al*., 2020). Other changes involve Asn99 (5.5 σ), Leu60 (with a difference density pair at ±4.0 σ), and Phe59 (4.2 σ and - 3.8 σ) shift by 0.5 – 1.0 Å, accompanying the rotation of the FMN on its axis. The distant residues located in the loop connecting Gβ and Hβ (Arg91, Asp93, Thr95, peaks above 4.0 σ) and adjacent to the C-terminal end of our construct are impacted (**Fig. 5c** **&** **Fig. 6c**), lending additional support to the changes in local protein dynamics around the C-terminal linker sequence implicated in signal propagation.

**Fig. 5.**
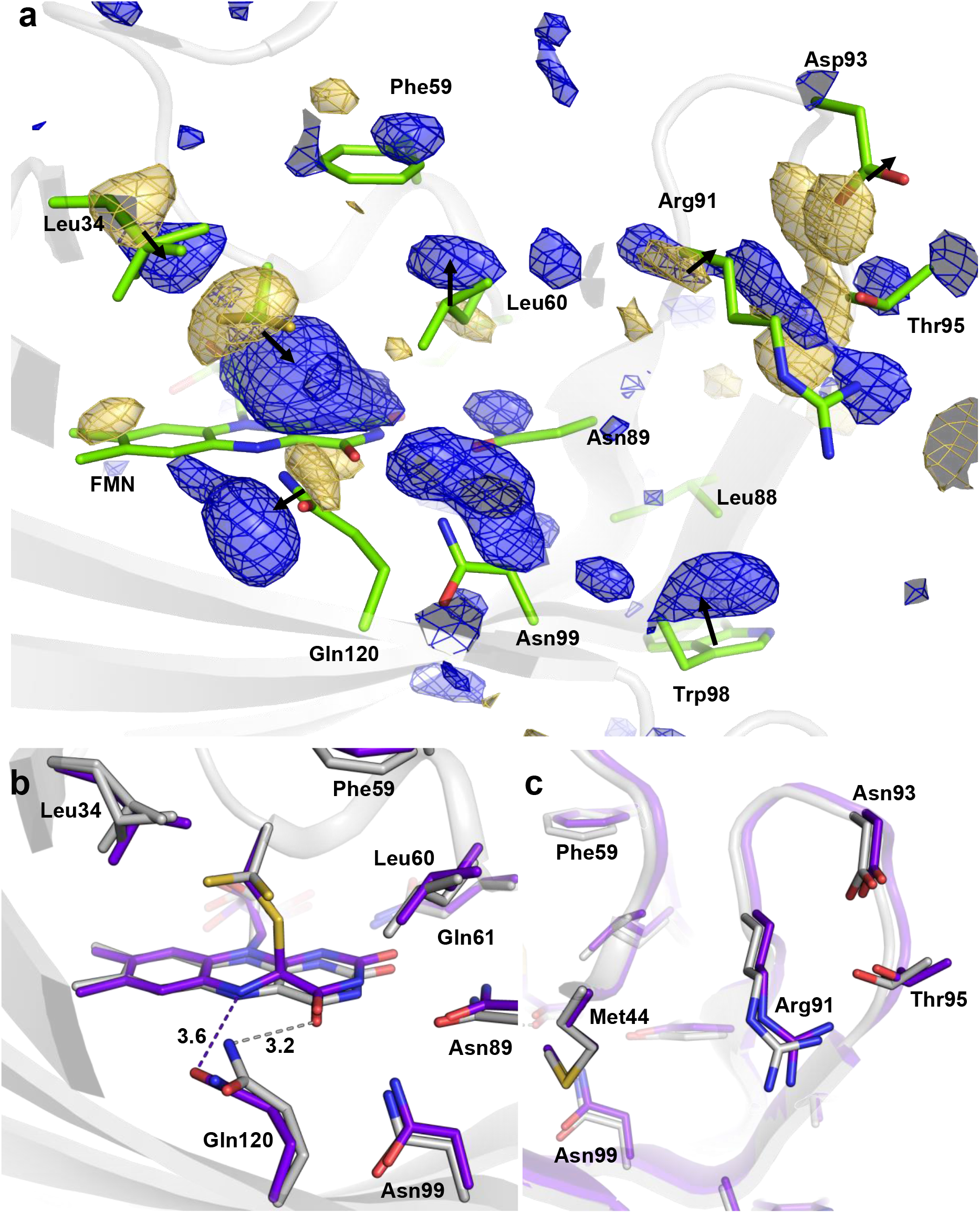
(**a**) *F*_obs_(2.5ms)-*F*_obs_(dark) maps contoured at 3 σ around the FMN chromophore and surrounding residues. Pairs of positive and negative peaks of density are indicated with an arrow. (**b**) Close-up superposition of the model coordinates of the refined light activated state (purple) and the dark state (grey) from SSX data on the flavin region, showing the rotation of Gln120 that is H-bonded to the protonated N_5_ of FMN. (**c**) Close-up view on the loop connecting β-strand H and I.

**Fig. 6.**
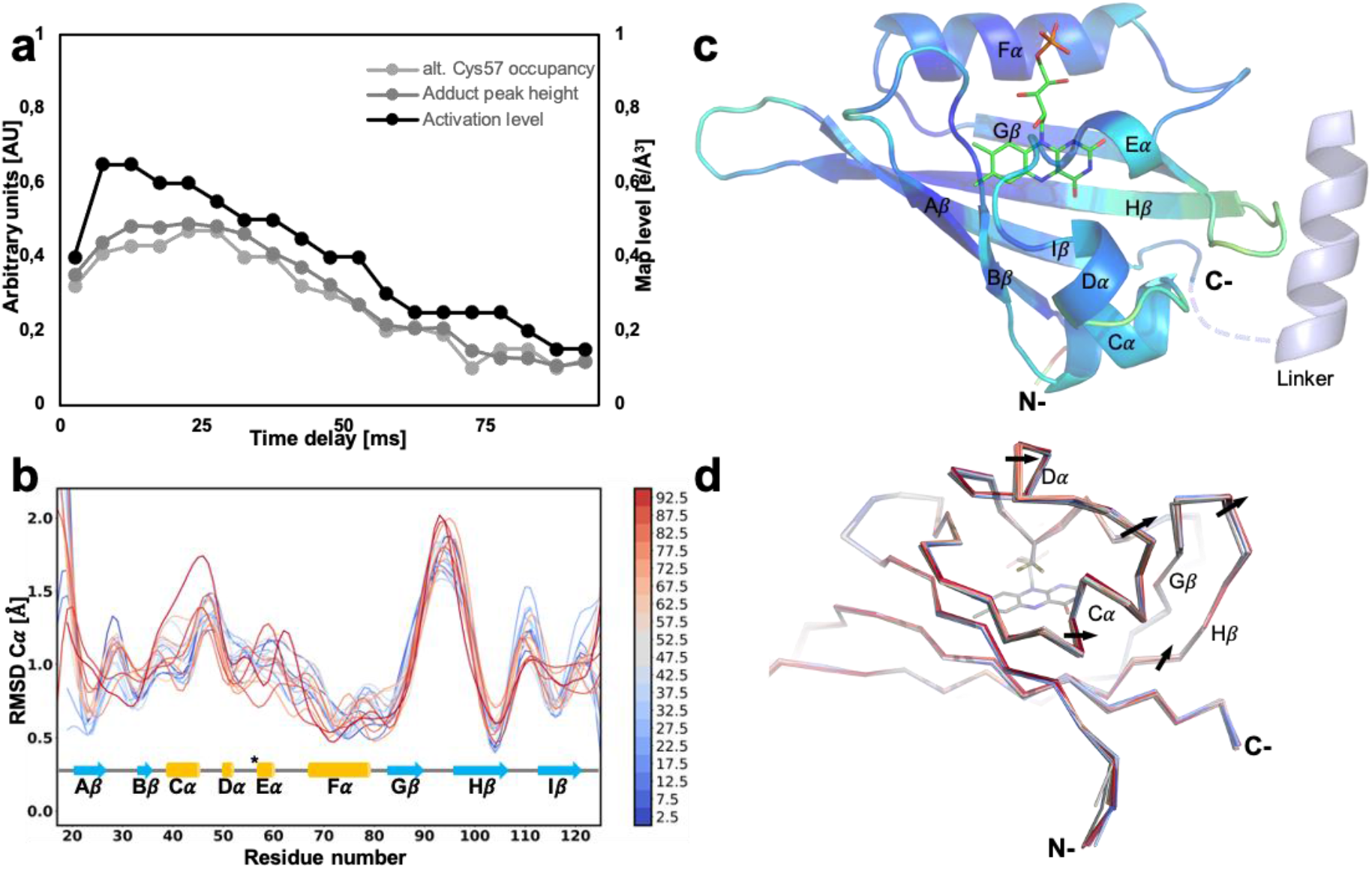
(**a**) Evolution of the activation level (black curve), the refined occupancy of Cys57 bound to FMN (light grey curve) and the height of the peak corresponding to the adduct in the*_F_* _light(n) – *F*_ ^dark^ map (grey) represented as a function of the time delay. (**b**) Evolution of the _obs obs_ root mean square deviation (RMSD) between the dark state model and the successive light states from 2.5 ms to 92.5 ms after photoactivation (blue to red curves) mapped over the secondary structure representation of *Cr*PhotLOV1. The reactive cysteine (Cys57) is indicated by a star. (**c**) Heat color cartoon representation of the average RMSD between the dark state and light states showing the secondary structures around the C-terminal part that are affected by the adduct formation. The C-terminal linker region with the LOV2 domain present in the full length phototropin is illustrated with grey dashes connecting an α-helix. (**d**) Superposition of light states from 2.5 (blue) to 92.5 ms after photoactivation (red) over the dark state (grey) with black arrows indicating the directionality of the structural change.

Subsequent time delays (*i.e.*, Δ*t* = 7.5 ms and 12.5 ms) initially display an increase in the strength of difference map peaks (such as the peak located on the covalent adduct, which reaches a maximum at Δ*t* = 22.5 ms), followed by a gradual decrease until all peaks (except for the peak on the covalent adduct) fall below ±3 σ at 82.5 ms (**Fig. 6a**, **Supplementary Fig. S6a** and **Fig. S6b**). This behavior aligns with the occupancy refinement results of the three alternate conformations of Cys57 (i.e., the two conformations from the dark state and the adduct) against the raw light datasets (refined without extrapolating structure factor amplitudes), which revealed an increase in the occupancy of the cysteinyl-FMN adduct alternate conformation up to Δ*t* = 22.5 ms, followed by a decrease over time. Furthermore, the trend is similar to the inferred activation levels (**Fig. S6a**). The initial increase in the active state signal and populations until Δ*t* = 22.5 ms likely results from a slight offset between the pump pulse and the X-ray interaction region. The decline in activation level likely results from the displacement of the continuously flowing stream section containing photoactivated crystals relative to the region probed by the X-ray beam. Indeed, at Δ*t* = 82.5 ms, the continuous sample stream has moved 51 μm since the Δ*t* = 0 (**Fig. S6&S5**). As a result, crystals probed by an X-ray beam at that time delay received less pump light (assuming a Gaussian distribution of pump pulse intensity). Despite the reduction in activation levels and signal intensity, the structural models could be refined against the extrapolated data up to 92.5 ms post-photoactivation (refinement statistics are presented in **Table 1**).

As anticipated, considering the time constant in the order of microseconds required for covalent bond formation, the most pronounced structural changes occur during the initial time delay (Δt = 2.5 ms). However, more subtle structural dynamics evolution can be observed by superimposing the dark state with subsequent light-activated states. Notably, Gβ-Hβ (0.7 Å at Δ*t* = 92.5 ms) and loop Hβ-Iβ (0.6 Å at Δ*t* = 32.5 ms) demonstrate significant divergence from the dark state, with the latter relaxing gradually back to the dark state conformation after Δ*t* = 32.5 ms (**Fig. 6b**). The structural motion of Gβ-Hβ appears to be primarily driven by the rotation of the FMN axis, pulling residues Asn89 and Asn99 along with it. Furthermore, while Leu101 does not display a fully rotated rotamer as observed for the homologous proteins, like photoreceptor *Pp*sB1-LOV from *Pseudomonas putida* or in other proteins where it is replaced by phenylalanine, such as *At*Phot2LOV2 from *Arabidopsis thaliana*, *Pt*Au1A (Aureochrome1A) from *Phaeodactylum tricornutum*, and Aureochrome 1 from *Vaucheria frigida* (see **Supplementary Table S1),** still a positive peak adjacent to this residue suggests about 15° rotation of the side chain. This rotation fills the space vacated by the twist of the flavin plane and the movement of Asn99. Intriguingly, this protein section flanks the N- and C-terminals connected to the LOV2 domain through a hinge region (although truncated in our construct; **Fig. 6c** and **6d**). Aumonier and colleagues (Aumonier et al., 2020) proposed that the rearrangement of Phe470 (in the case of *Cr*PhotLOV1, Leu101) impacts Leu456 (here, Leu87) and, by extension, the groove stabilizing the Jα linker helix. These observations collectively support a hypothesis that signal propagation in *Cr*PhotLOV1 is related to extended changes in local protein dynamics (Dittrich *et al*., 2005; Pfeifer *et al*., 2009), rather than a conformational change of a specific residue. Additionally, accumulating structural changes in Gβ-Hβ over time could promote LOV domain oligomerization, resulting in a long-lasting signaling state (Nakasone *et al*., 2018, 2019). This observation aligns with spectroscopic characterizations of full-length phototropin, demonstrating a time constant of 77 ms for helix structuration (Nakasone *et al*., 2018, 2019).

### Covalent adduct conformation in photoactivated states

To date, 103 structures of LOV domains have been deposited in the Protein Data Bank, with 22 corresponding to a photostationary light state (**Supplementary Table S1**). Two distinct conformations of the covalent adduct have been noted (**Fig. 7c**). The predominant adduct conformation across the deposited structures features the Cys57 cysteinyl group oriented similarly to conformation B of the resting state (**Fig. 7a**), as it forms a covalent bond with the FMN C4a in the *sp3* configuration. The alternative geometry, described in the seminal *Cr*PhotLOV1 paper (Fedorov *et al*., 2003), involves the entire Cys57 residue being translated by 1.4 Å and oriented in the opposite direction, closer to conformation A of Cys57 of the resting state (**Fig. 7c**). However, the conformation reported by Fedorov *et al*. of the reactive cysteine has not been observed in other photostationary states of homologous proteins obtained at high resolution (**Supplementary Table S1**). Additionally, the FMN isoalloxazine ring would need to move 1.1 Å toward the sulfur, with a twist of the pyrimidine side of the ring, which is not confirmed by our high-resolution room temperature crystallographic data. In the present work, the models of the photoactivated states exhibit far better fit when the more common adduct geometry, i.e., closer to conformation B of Cys57 of the resting state, is employed (**Fig. 7b**). Thus, contrasting with the adduct geometry originally determined (**Fig. 7c**). Resolved in this work the cysteinyl-FMN adduct conformation should have significant implications for subsequent molecular dynamics and QM/MM calculations aimed at understanding activation and signaling in LOV photoproteins.

**Fig. 7.**
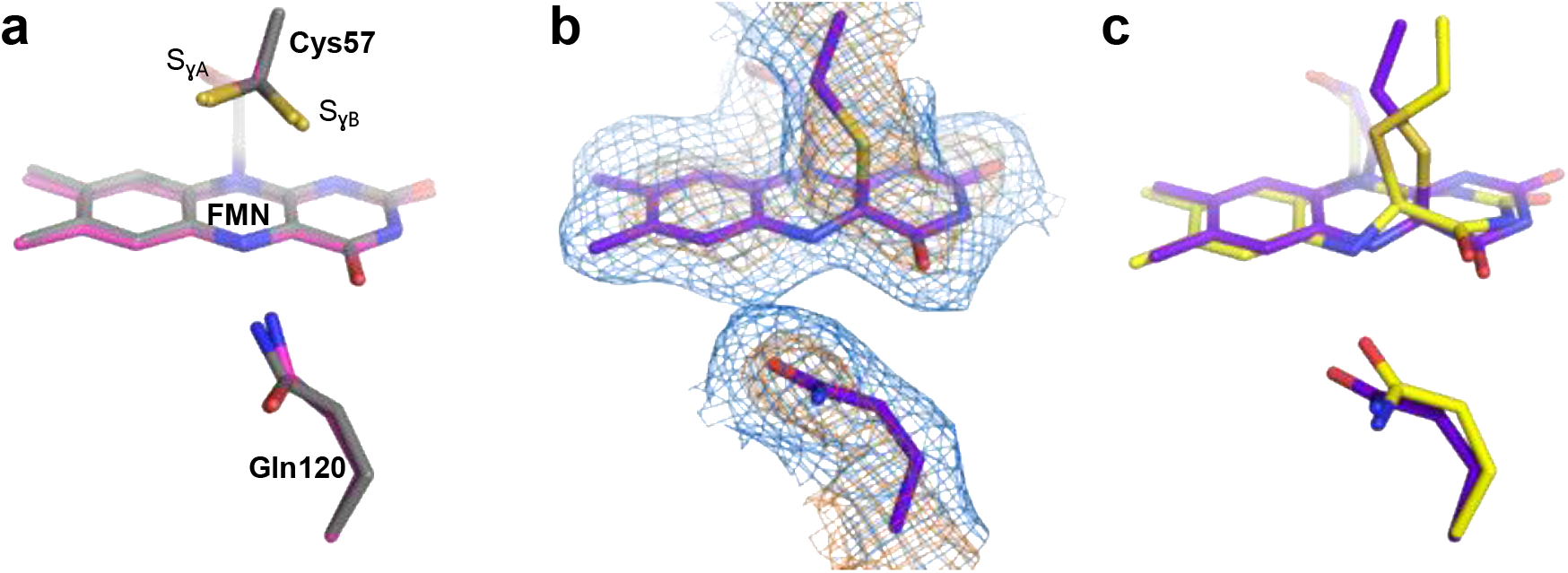
**Comparison of dark and active state models of *Cr***Phot**LOV1 with previously published structures.** (**a**) Superposition of dark state model from Fedorov *et al*., 2003 obtained at cryogenic temperature (magenta) with our dark state (gray) obtained at room temperature. (**b**) 2*F* _light(n) – *F*_ extrapolated electron density maps at Δ*t* = 2.5 ms shown at _obs calc_ 1.0 and 3.0 σ (blue and orange mesh, respectively) around the Cys57-FMN and Gln120. (**c**) Superposition of the light-adapted state from Fedorov *et al*., 2003 (yellow) with our 2.5 ms structure (purple) showing the translation of Gln120 and the difference in the geometry of the FMN-Cys57 adduct with the original 1N9L structure. Structural coordinates were superposed within *Pymol* with the *cealign* algorithm.

## Conclusion

The advancements in brighter synchrotron beams and high-frame-rate low-noise photon-counting X-ray detectors have rekindled interest in obtaining protein structures under near-native room temperature conditions (Stellato *et al*., 2014; Owen *et al*., 2014; Fischer, 2021). Moreover, technology transfer from X-ray Free Electron Lasers (XFELs) to synchrotron beamlines, such as sample delivery instrumentation, has led to a growing number of studies focused on probing the structural dynamics of proteins on millisecond to second timescales at synchrotron light sources (Martin-Garcia, 2021).

In this work, we presented a TR-SSX experiment on *Cr*PhotLOV1, along with the protocol and its optimization for producing the microcrystals required. This protocol, which identified HEC as an optimal carrier matrix, facilitates the collection of TR-SSX data and could be readily adapted for studying other soluble proteins using a similar approach. Prior to crystallographic studies in crystallo spectroscopy was employed to assess protein photoreactivity. In the following pump-probe experiment, we captured snapshots of the photoactivated state from Δ*t* = 2.5 ms to 92.5 ms at a time resolution of 5 ms, which is an order of magnitude faster than previous works on *At*Phot2LOV2 (Aumonier *et al*., 2020). These data offer new insights into the fine changes of the LOV1 domain occurring in the millisecond time range, correlating with spectroscopic signal propagation studies. Furthermore, supported by the high-resolution crystallographic data, we resolve the geometry of the *Cr*PhotLOV1 thioadduct formed upon photoactivation, a controversial topic based on the previous reports. This study detailing steps from sample optimization to data analysis can collectively serve as a framework for routine time-resolved crystallography at synchrotrons.

## Acknowledgements

We thank the staff of the X06SA and X10SA beamlines at the Swiss Light Source of the Paul Scherrer Institute for their assistance. G. Gotthard is acknowledging the PSI-FELLOW-II-3i programme from the Paul Scherrer Institute.

## Supplementary materials

### Supplementary Tables

**Table S1.**
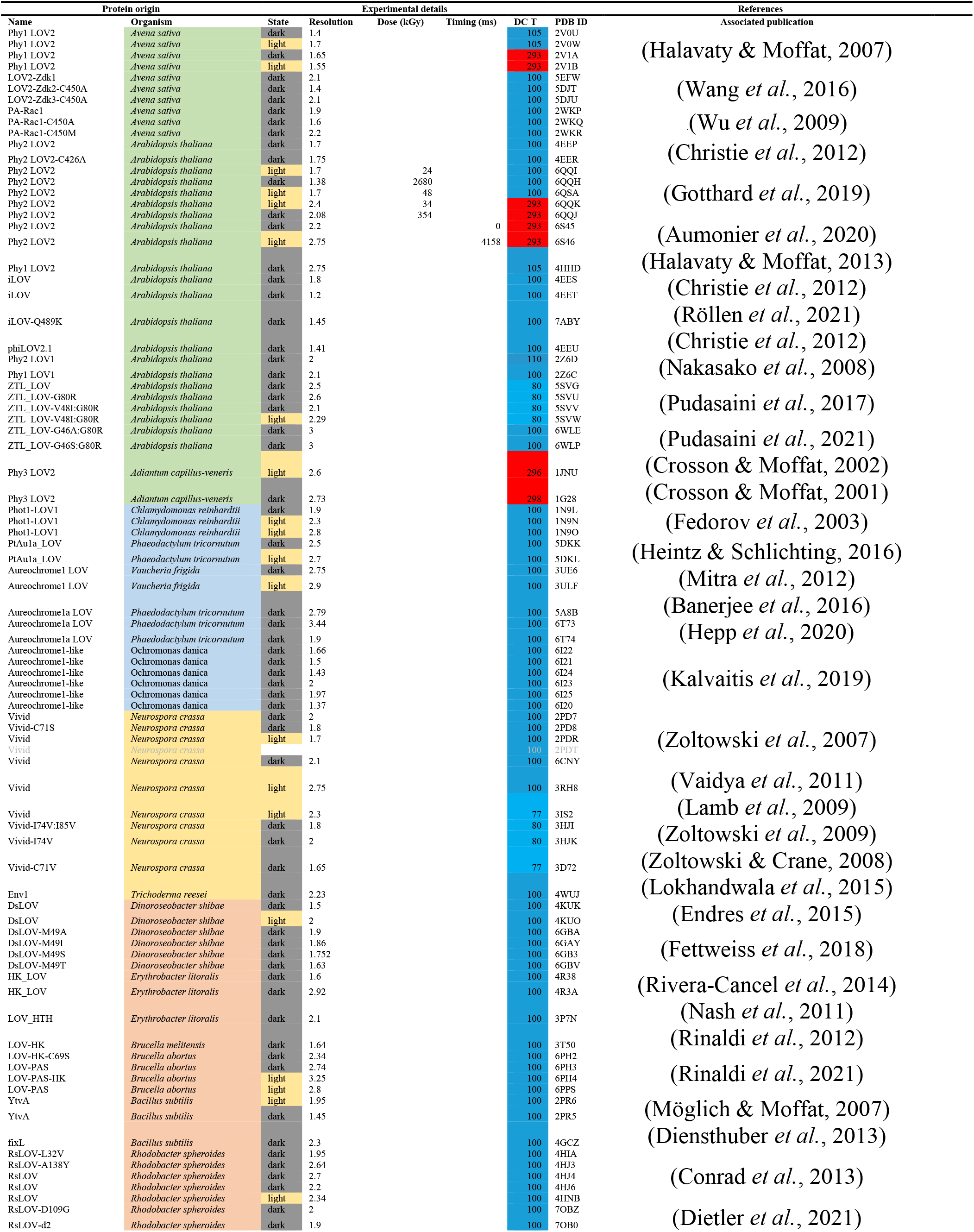
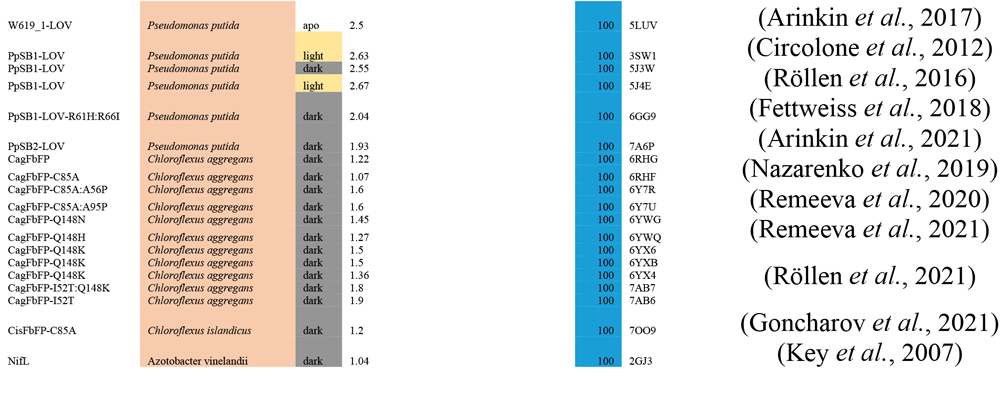
Structures of LOV domains in the Protein Data Bank.

### Supplementary Figures

**Fig. S1.**
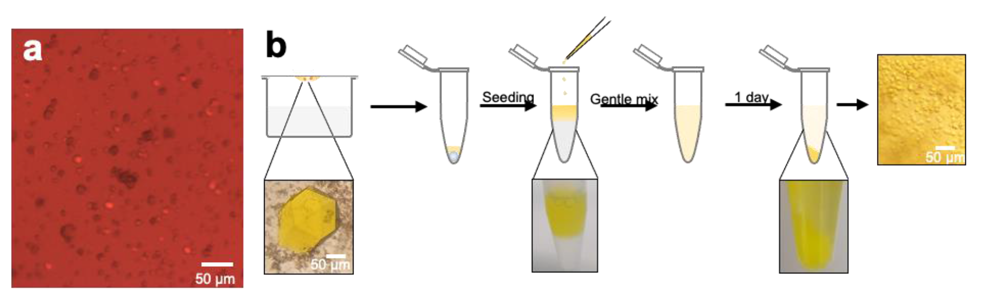
Crystallization and preparation of *Cr*PhotLOV1 microcrystals. (**a**) Microcrystals obtained in crystallization screening observed under the microscope with a red filter after one day. (**b**) Protocol for preparation of *Cr*PhotLOV1 microcrystals with the batch crystallization method, *i*) macrocrystals are grown with the hanging drop crystallization method, *ii*) crystal seeds are then prepared from macrocrystals with Hampton research seeding beads and *iii*) are mixed with protein solution which is added dropwise into an Eppendorf tube containing the crystallization condition, *iv*) after a gentle mixing by inverting the tube several times, and *v*) one day at room temperature, *vi*) microcrystals appears and sediment at the bottom of the tube.

**Fig. S2.**
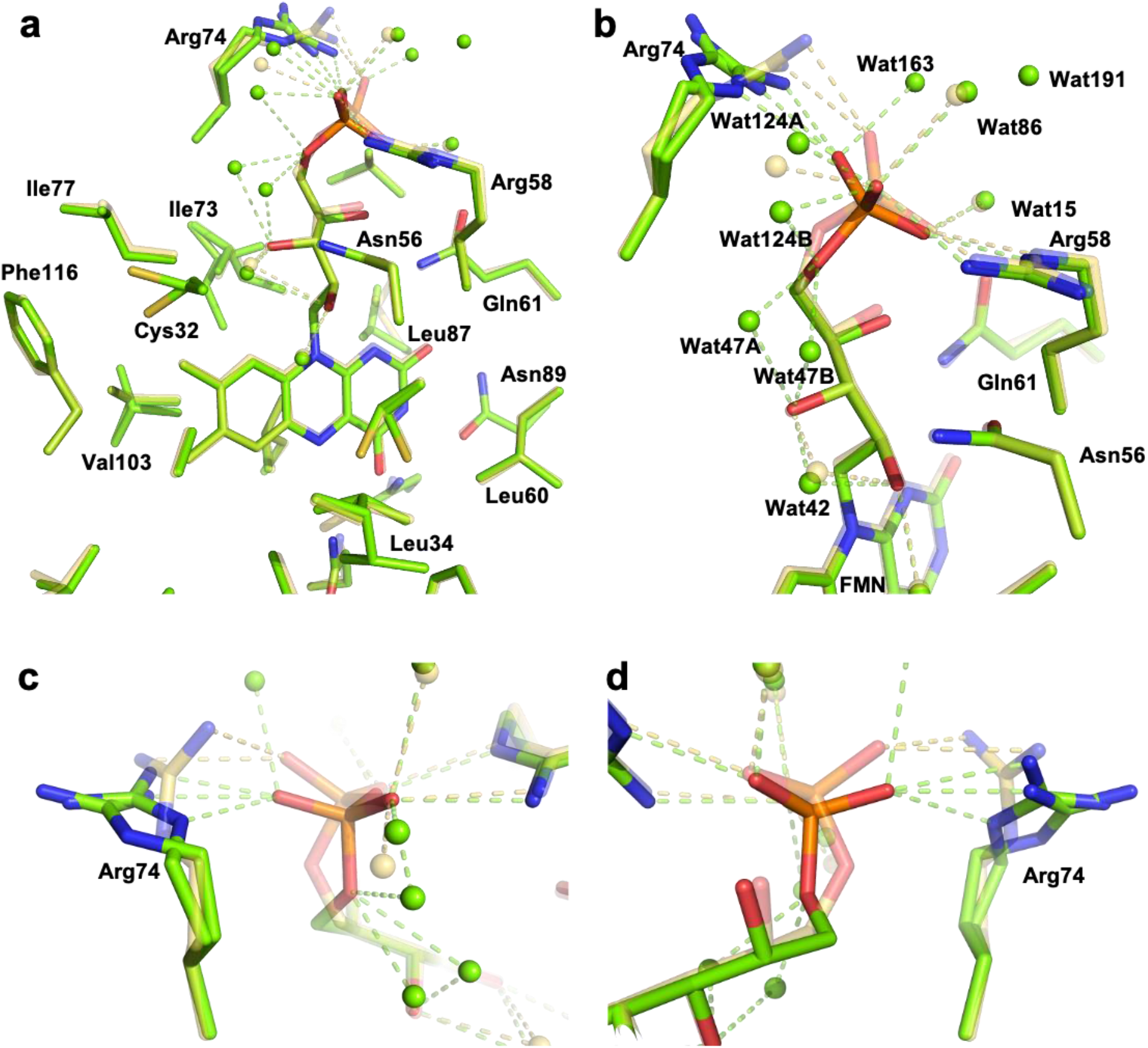
Differences in the coordination net around the FMN in *Cr*PhotLOV1 dark-state structures at cryogenic temperature. The structure solved in this work (green) is superimposed on the structure 1N9L (transparent yellow) (**a**) far view of the FMN environment, (b) close up on the phosphoribityl tail, (c) and (d) close-up view on the Arg74.

**Fig. S3.**
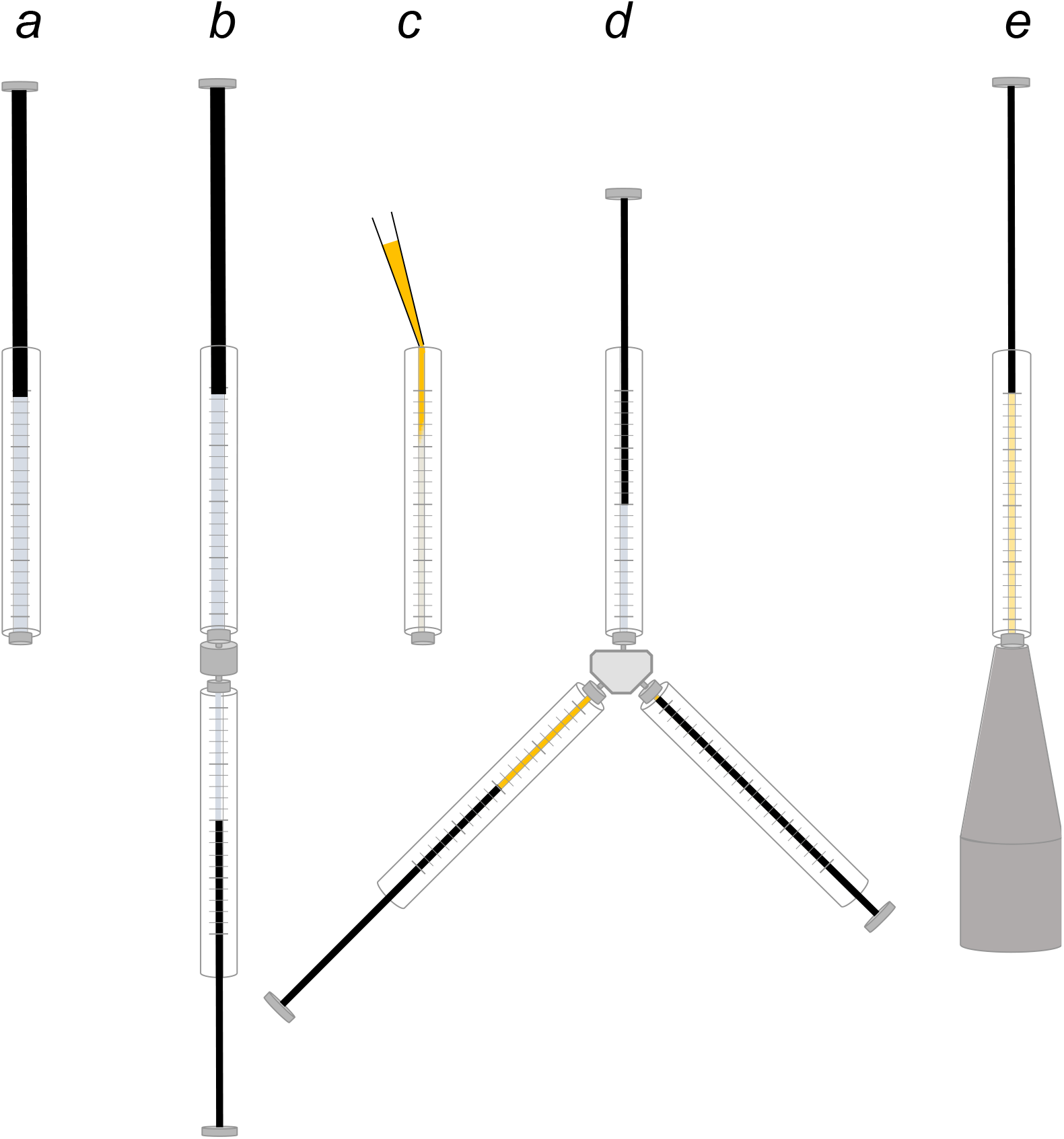
Protocol for embedding *Cr*PhotLOV1 crystals in HEC. (**a**) HEC is rehydrated with crystallization buffer in 500 ml Hamilton syringe. (**b**) The required amount of cellulose is transferred in a 100 ml Hamilton syringe. (**c**) LOV crystals in solution are inserted at the back of a 100 ml Hamilton syringe with caution to avoid bubbles. (**d**) LOV crystals are mixed with the cellulose using a three-way coupler until homogeneity. (**e**) Cellulose embedded crystals are then loaded in the reservoir of the HVE injector.

**Fig. S4.**
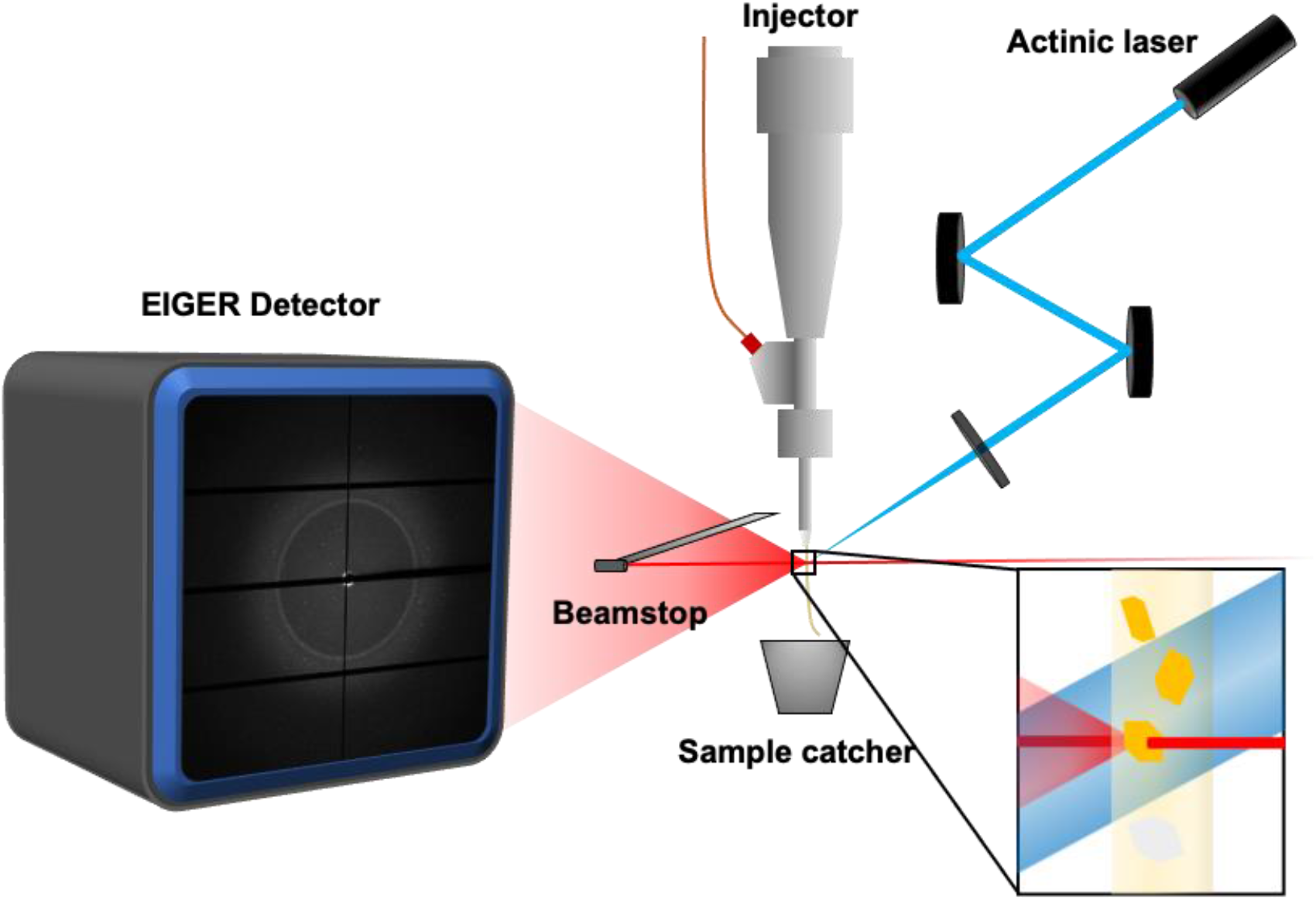
Time resolved serial synchrotron crystallography setup. LOV crystals are serially injected onto the path of the X-ray synchrotron beam. Crystals are photoexcited using a 470 nm focused laser that is synchronized with the trigger of the detector. Diffraction patterns are collected following the defined data collection scheme (**Fig. S5**).

**Fig. S5.**
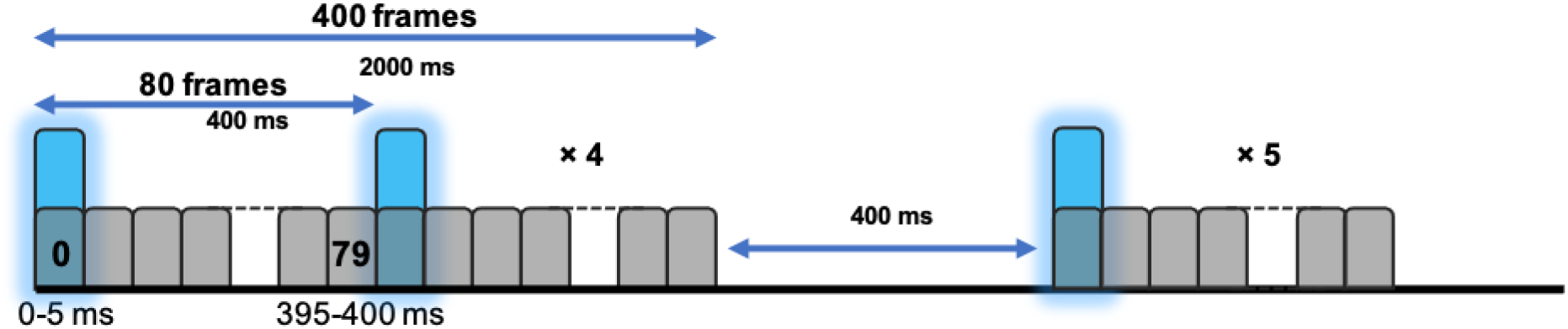
Schematic representation of the pulse / detector organization for the TR-SSX data collection on LOV1. The activation sequence is composed of one 5 ms frame collected with the laser diode on (blue histogram), followed by 79 frames collected without illumination (gray histogram). The sequence is repeated 5 times, after which one activation sequence is skipped.

**Fig. S6.**
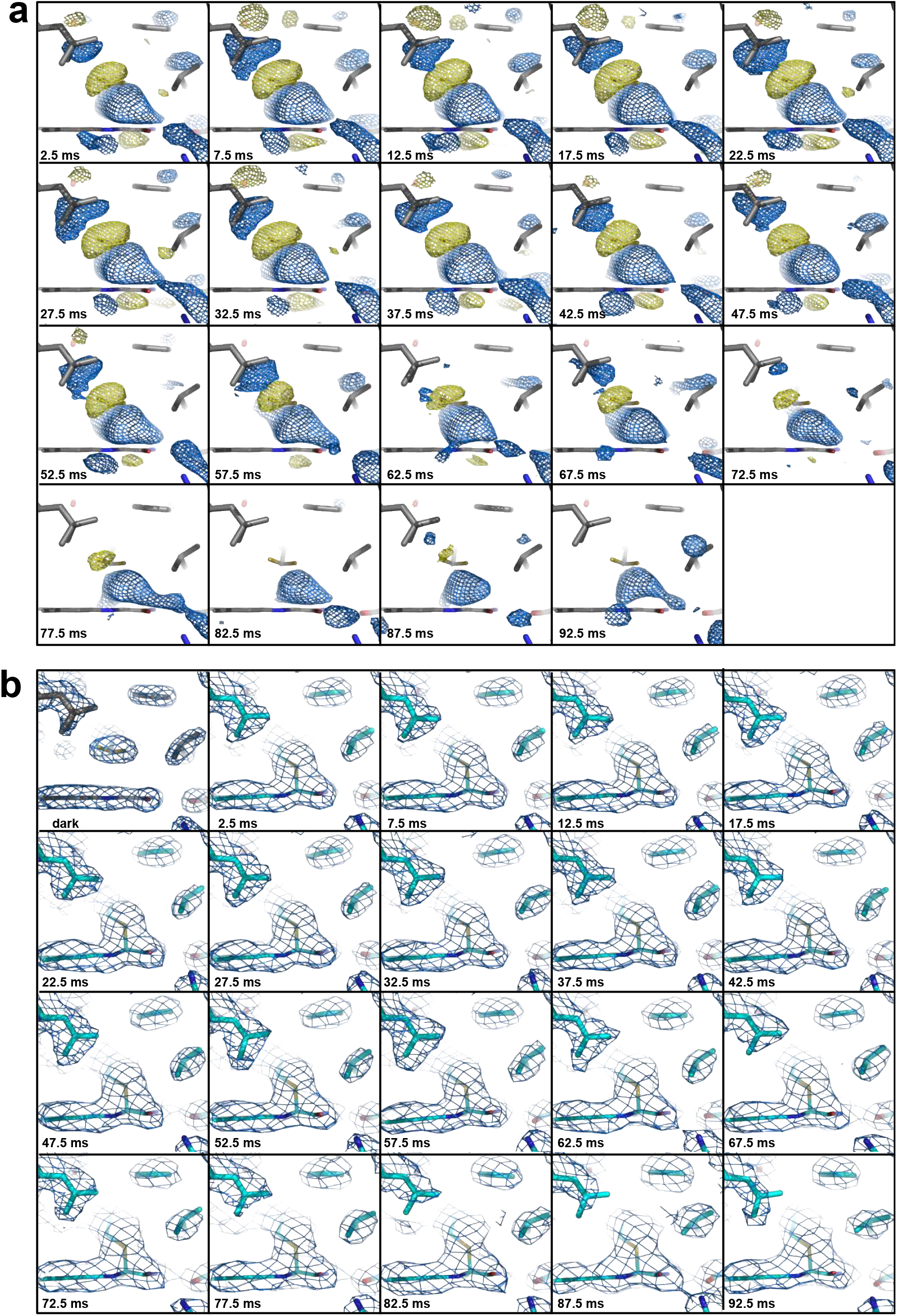
Fourier difference and extrapolated electron density maps. (**a**) *F*_obslight(n)_ – *F*_obsdark_ from 2.5 to 92.5 ms after photoactivation surrounding the adduct represented at ±3.0 σ showing the slow decrease signal in the maps. (**b**) Dark state 2*F*_obs_ – *F*_calc_ and 2*F*_ext_ – *F*_calc_ extrapolated maps from 2.5 to 92.5 ms.

